# Murine cytomegaloviruses m139 targets DDX3 to curtail interferon production and promote viral replication

**DOI:** 10.1101/2020.04.17.046516

**Authors:** Olha Puhach, Eleonore Ostermann, Christoph Krisp, Giada Frascaroli, Hartmut Schlüter, Melanie M. Brinkmann, Wolfram Brune

## Abstract

Cytomegaloviruses (CMV) infect many different cell types and tissues in their respective hosts. Monocytes and macrophages play an important role in CMV dissemination from the site of infection to target organs. Moreover, macrophages are specialized in pathogen sensing and respond to infection by secreting cytokines and interferons. In murine cytomegalovirus (MCMV), a model for human cytomegalovirus, several genes required for efficient replication in macrophages have been identified, but their specific functions remain poorly understood. Here we show that MCMV m139, a gene of the conserved US22 gene family, encodes a protein that interacts with the DEAD box helicase DDX3, a protein involved in pathogen sensing and interferon (IFN) induction, and the E3 ubiquitin ligase UBR5. DDX3 and UBR5 also participate in the transcription, processing, and translation of a subset of cellular mRNAs. We show that m139 inhibits DDX3-mediated IFN-β induction and is necessary for efficient viral replication in bone-marrow derived macrophages. In vivo, m139 is crucial for viral dissemination to local lymph nodes and to the salivary glands. An m139-deficient MCMV also replicated to lower titers in SVEC4-10 endothelial cells. This replication defect was not accompanied by increased IFN-β transcription, but was rescued by knockout of either DDX3 or UBR5. Moreover, m139 co-localized with DDX3 and UBR5 in viral replication compartments in the cell nucleus. These results suggest that m139 inhibits DDX3-mediated IFN-β production in macrophages and antagonizes DDX3 and UBR5-dependent functions related to RNA metabolism in endothelial cells.

**Author Summary:** Human cytomegalovirus is an opportunistic pathogen that causes severe infections in immunocompromised individuals. The virus infects certain types, such as macrophages and endothelial cells, to ensure its dissemination within the body. Little is known about the viral factors that promote a productive infection of these cell types. The identification of critical viral factors and the molecular pathways they target can lead to the development of novel antiviral treatment strategies. Using the mouse cytomegalovirus as a model, we studied the viral m139 gene, which is important for virus replication in macrophages and endothelial cells and for dissemination in the mouse. This gene encodes a protein that interacts with the host proteins DDX3 and UBR5. Both proteins are involved in gene expression, and the RNA helicase DDX3 also participates in mounting an innate antiviral response. By interacting with DDX3 and UBR5, m139 ensures efficient viral replication in endothelial cells. Importantly, we identify m139 as a new viral DDX3 inhibitor, which curtails the production of interferon in macrophages.

## Introduction

Cytomegaloviruses (CMVs) have a broad tissue tropism and infect a wide variety of cells types. Cells of the myeloid lineage play an important role in CMV dissemination and pathogenesis: latently infected monocytes disseminate the virus to target organs [1,2]. Differentiation of monocytes into macrophages as they extravasate into tissues triggers the lytic replication cycle and the production of infectious virus, which transmits the infection to the surrounding parenchymal cells. Efficient replication in macrophages requires that the virus is able to curtail its recognition by the innate immune system [3]. This is particularly important in macrophages, as these cells express numerous sensors that activate host antiviral defenses and induce the production of cytokines and interferons [4].

Another cell type involved in CMV dissemination and pathogenesis are endothelial cells. They support productive virus replication and are thought to contribute to dissemination via the blood stream [5]. Moreover, endothelial cells have been proposed to be sites of latency [6].

Murine cytomegalovirus (MCMV) serves as a small animal model for human cytomegalovirus (HCMV). It encodes a number of genes known to be important for efficient replication in macrophages, several of which belong to the US22 gene family. This gene family comprises 12 members in both, HCMV and MCMV, that contain up to four conserved sequence motifs [7,8]. The functions of most US22 family genes remain unknown; only in a few cases has the mechanism of action been resolved. MCMV genes m142 and m143 are the only US22 family genes essential for viral replication [8]. The m142 and m143 proteins form a heterodimeric dsRNA-binding complex and inhibit the activation of the dsRNA-dependent protein kinase, PKR [9,10]. Their homologs in HCMV, TRS1 and IRS1, fulfill essentially the same function [11,12]. Inhibition of PKR is essential for CMV replication in cell culture and in vivo [13-16]. Another well-characterized US22 family gene is MCMV M36 and its homolog in HCMV, UL36. The M36 and UL36 proteins prevent apoptosis by inhibiting caspase-8 activation [8,17]. This function is particularly important for virus replication in macrophages and dissemination in vivo [18-20].

Besides M36, the MCMV genes m139, m140, and m141 promote efficient replication of the virus in macrophages [8,21-23]. The three genes are clustered within the MCMV genome and their protein products can bind to each other, suggesting that they might function cooperatively [24]. On the other hand, the three proteins appear to have individual functions. The m140 protein is required for efficient capsid assembly in macrophages [25]. However, the underlying mechanism remains incompletely understood. The m140 protein interacts with m141 and stabilizes it by protecting it from proteasomal degradation [26]. The m141 protein directs m140 to a perinuclear region of infected macrophages adjacent to the microtubule organizing center, suggesting an involvement of aggresomes [25].

The least is known about m139. Its requirement for efficient replication in IC-21 macrophages has been documented [8], but m139 was not required for the functions described for m140 and m141 [25,26]. However, a recently performed screen for MCMV inhibitors of interferon-β (IFN-β) induction identified m139 as a candidate [27]. These findings prompted us to investigate the function of m139, its role for viral replication in macrophages and other cell types, and its requirement for viral dissemination in vivo.

In this study, we show that m139 is required for efficient virus replication in bone marrow-derived macrophages and endothelial cells in vitro and for viral dissemination in vivo. By affinity purification and mass spectrometry, we identified the host proteins DDX3 and UBR5 as interactors of m139. Knockouts of DDX3 or UBR5 in endothelial cells rescued the replication defect of an m139-deficient MCMV. Moreover, m139 inhibited DDX3-dependent IFN-β induction in macrophages to facilitate efficient virus replication in these cells. This study reveals that m139 counteracts the host factors DDX3 and UBR5 to subvert host antiviral responses and promote viral replication.

## Results

### The MCMV protein m139 is expressed with early kinetics and recruited to viral replication compartments

The m139 ORF is transcribed with early kinetics and encodes two proteins of 72 and 61 kDa [21]. To further characterize the m139 proteins in the absence of a specific antibody, we generated a recombinant MCMV expressing a C-terminally HA-tagged m139. By BAC recombineering an HA tag sequence was inserted at the 3’ end of the m139 ORF. We verified the classification of m139 as an early protein by using a cycloheximide (CHX) release assay. Fibroblasts were infected in the presence of CHX, a translation inhibitor. Four hours post infection, cells were washed and incubated either with normal medium, in order to relieve the inhibition, or with medium containing actinomycin D (ActD), a transcription inhibitor, to selectively allow the translation of viral immediate-early transcripts. As shown in Figure 1A, m139 was expressed in untreated cells and after release from the CHX block, but not in the presence of ActD. Thus, m139 is expressed with early kinetics, similar to the M112-113 encoded Early 1 (E1) proteins.

**Figure 1.**
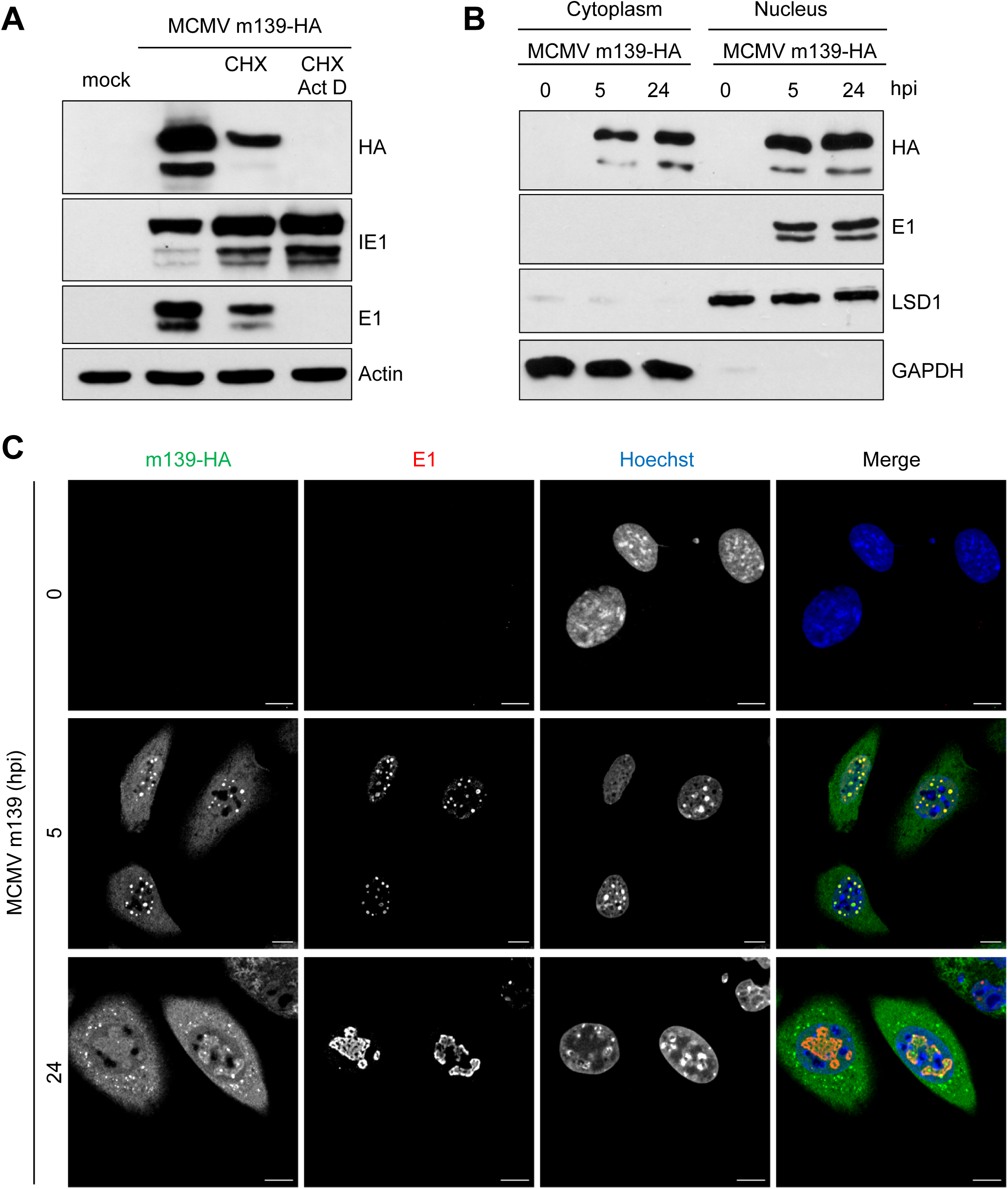
MCMV m139 is expressed with early kinetics and localizes to the the cytoplasm and viral replication compartments in the nucleus. (A) 10.1 fibroblasts were infected with MCMV m139-HA (MOI=3) in the presence of CHX (5 μg/ml) or left untreated. Four hpi, the CHX-containing medium was replaced by normal medium or medium containing ActD (50 μg/ml). After 4 hours, viral protein expression was analyzed by immunoblotting. (B) 10.1 fibroblasts were infected with MCMV m139-HA (MOI=5). Cell lysates were harvested at 5 and 24 hpi, separated into nuclear and cytoplasmic fractions, and analyzed by immunoblotting. (C) 10.1 fibroblast were infected with MCMV m139-HA (MOI=1), fixed at 5 and 24 hpi, and subjected to immunofluorescence. Mock-infected cells were used as a control. m139 was detected with an anti-HA antibody and the viral E1 (encoded by M112-113) proteins with an E1-specific antibody. Nuclei were counterstained with Hoechst 33342. Representative images taken by confocal microscopy are shown. Scale bar, 10 μm.

To clarify the subcellular localization of m139 during infection, we used a cell fractionation assay to separate the nuclear from the cytoplasmic fraction. In fibroblasts infected with MCMV m139-HA, the m139 protein was found in both nuclear and cytoplasmic fractions at early and late times post infection (Figure 1B), consistent with a previous study [21]. By immunofluorescence, m139 was also detected in the cytoplasm and in the nucleus. It was found dispersed throughout the cytoplasm at early times and was enriched in perinuclear dots at late times post infection (Figure 1C). In the nucleus, m139 co-localized with the viral E1 proteins (Figure 1C). The MCMV E1 proteins are encoded by the M112-113 gene, exist in four isoforms [28,29], and are markers for viral replication compartments [30].

An in silico analysis of the m139 amino acid sequence revealed the presence of a nuclear export signal (NES). The predicted NES sequence SEIRVLRGVDLSD was mapped to residues 215-227. Consistent with this prediction, the m139 protein was detected exclusively in the cytoplasm of cells upon ectopic expression of m139 (Figure S1A). However, co-transfection of an E1 expression vector was sufficient to cause m139 accumulation in E1-positive dots within the nucleus (Figure S1B), suggesting that the viral E1 proteins recruit m139 to pre-replication compartments.

### m139 is required for efficient MCMV replication in macrophages and endothelial cells

A previous study has reported that m139 is required for MCMV replication to high titers in IC-21 peritoneal macrophages [8]. In order to investigate the role of m139 for MCMV replication in different cell types of mouse origin, we constructed an MCMV m139 knockout mutant by introducing a stop at codon position 32 of the m139 ORF. The genome integrity of the MCMV m139-HA and m139*stop* mutants were verified by deep sequencing to rule out accidental mutations elsewhere in the genome. We compared the replication properties of the MCMV m139*stop* mutant to the wildtype MCMV m139-HA by multistep replication kinetics in 10.1 fibroblasts, TCMK-1 epithelial cells, SVEC4-10 endothelial cells, and immortalized bone marrow-derived macrophages (iBMDM). In 10.1 fibroblasts and TCMK-1 epithelial cells, the MCMV m139*stop* mutant replicated to titers comparable to MCMV m139-HA (Figure 2A and B). By contrast, MCMV m139*stop* replicated to substantially lower levels in SVEC4-10 endothelial cells (Figure 2C) and iBMDM (Figure 2D). These results demonstrated that m139 is required for efficient MCMV replication in specific cell types such as macrophages and endothelial cells, but is dispensable in fibroblasts and epithelial cells. MCMV m139-HA and WT MCMV replicated to similar titers in these cell types, indicating that the C-terminal HA tag did not negatively affect MCMV replication (Figure 2C, D).

**Figure 2.**
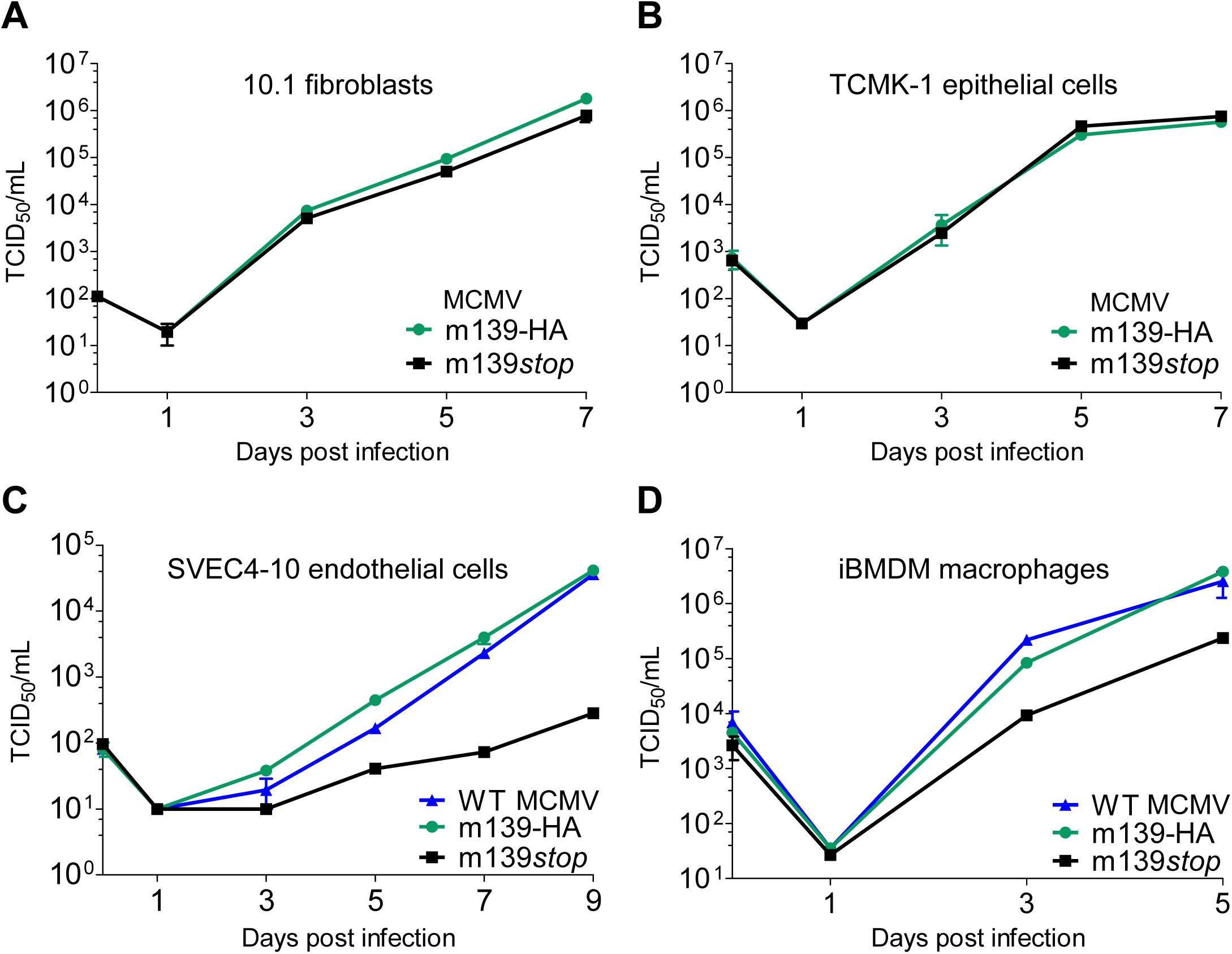
MCMV m139 is important for viral replication in macrophages and endothelial cells. Multistep replication kinetics of WT MCMV, MCMV m139-HA and MCMV m139*stop* in murine 10.1 fibroblasts (A), TCMK-1 epithelial cells (B), SVEC4-10 endothelial cells (C), and immortalized bone marrow-derived macrophages (iBMDM) (D). Cells were infected at an MOI of 0.01. Virus released by infected cells into the supernatant was quantified by titration. Viral titers are shown as means ±SEM of triplicates.

### m139 interacts with host proteins DDX3 and UBR5

In order to gain further insight into the role of m139 in MCMV replication, we wanted to identify m139-interacting proteins in MCMV-infected cells. We used stable isotope labeling of amino acids in cell culture (SILAC [31]) combined with affinity purification and mass spectrometry (AP-MS) to identify viral and host proteins interacting with m139. SVEC4-10 cells were infected with MCMV m139-HA or WT MCMV, and m139 was immunoprecipitated using an anti-HA affinity matrix. Proteins identified with less than 4-fold enrichment in the MCMV m139-HA sample (log_2_ ratio <2) or less than 2 unique peptides detected were excluded from further analysis. Using these criteria, 11 putative interaction partners of m139 were identified (Table 1). The candidate list included MCMV proteins m140 and m141, which are known interaction partners of m139 [24]. Among the host proteins, we focused on those previously reported to affect CMV replication or type I IFN signaling: the ATP-dependent RNA helicase DDX3 (a.k.a. DEAD box protein 3), the E3 ubiquitin-protein ligase UBR5 (a.k.a. EDD1), and the interferon-induced protein with tetratricopeptide repeats 1 (IFIT1) [32-36]. To verify the interactions of m139 with DDX3, UBR5, and IFIT1, m139-HA or M45-HA as control were immunoprecipitated from lysates of cells infected with MCMV m139-HA or MCMV M45-HA. As shown in Figure 3A, DDX3 and UBR5 co-precipitated with m139, but IFIT1 did not. None of the three cellular proteins co-precipitated with M45. Interestingly, the viral E1 proteins did not co-precipitate with m139, even though the E1 proteins mediate the recruitment of m139 to nuclear replication compartments (Figures S1). These finding suggested that the viral m139 and E1 proteins interact indirectly or through a weak interaction.

**Table 1.** Proteins interacting with m139 identified AP-MS.

| Protein name | No. of peptides in 1<sup>st</sup> replicate | Ratio L/H | No. of peptides in 2<sup>nd</sup> replicate | Ratio H/L |
| --- | --- | --- | --- | --- |
| Myosin phosphatase Rho-interacting protein (Mrip) | 14 | 56.3 | 11 | 53.1 |
| MCMV protein m141 | 31 | 46.7 | 27 | 58.7 |
| MCMV protein m140 | 26 | 39.5 | 19 | 35.9 |
| E3 ubiquitin-protein ligase UBR5 | 12 | 14.8 | 8 | 31.3 |
| Ankycorbin | 20 | 18.9 | 12 | 22.5 |
| GEM-interacting protein (Gmip) | 8 | 16.4 | 5 | 20.3 |
| Interferon-induced protein with tetratricopeptide repeats 1 (IFIT1) | 13 | 7.1 | 12 | 14.7 |
| ATP-dependent RNA helicase DDX3 | 2 | 7.9 | 2 | 5.9 |
| Codanin-1 (Cdan1) | 3 | 3.7 | 3 | 7.8 |
| MCMV protein m142 | 9 | 3.1 | 12 | 2.6 |
| Unconventional myosin-Ic (Myo1c) | 2 | 2.0 | 2 | 2.3 |

**Figure 3.**
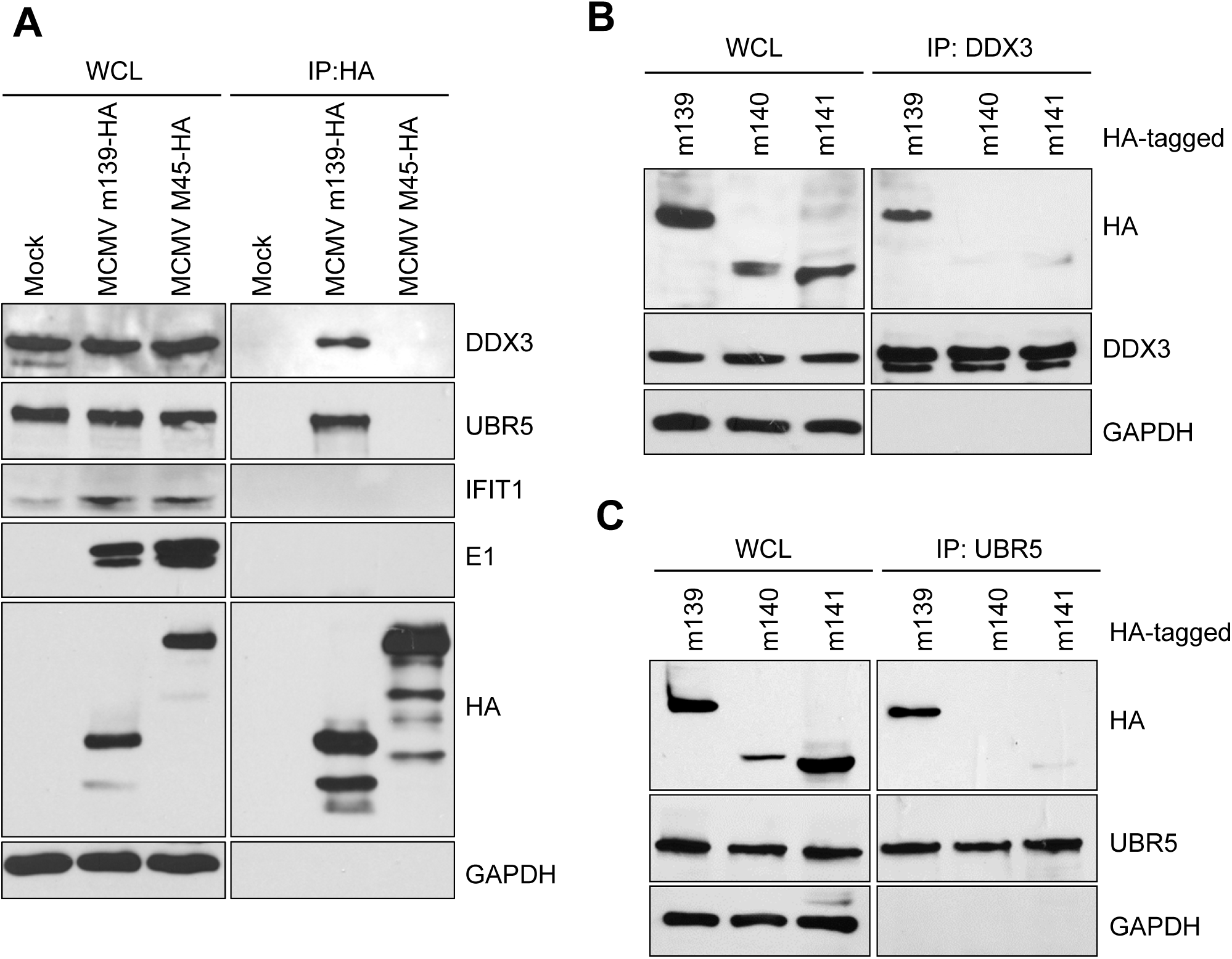
m139 interacts with DDX3 and UBR5. (A) SVEC4-10 cells were infected with MCMV m139-HA or M45-HA at an MOI of 5. Cell lysates were collected 24 hpi and subjected to immunoprecipitation (IP) using an anti-HA affinity matrix. Co-precipitating proteins were detected by immunoblotting with specific antibodies. (B, C) HEK-293A cells were transfected with expression plasmids encoding HA-tagged MCMV proteins m139, m140, or m141. Cell lysates were subjected to immunoprecipitation (IP) using anti-DDX3 (B) or anti-UBR5 (C) antibodies. Co-precipitating proteins were detected by immunoblotting as in (A).

DDX3 and UBR5 are multifunctional host proteins known to participate in fundamental cellular processes such as transcription, translation, and cell proliferation [37,38]. The RNA helicase DDX3 exists in two isoforms, DDX3X and DDX3Y, which are thought to have redundant functions. While DDX3Y expression is largely confined to the male germ line, DDX3X (hereafter referred to as DDX3) is ubiquitously expressed [39]. DDX3 has multiple functions in the context of RNA metabolism due to its ability to reorganize RNA secondary structures and ribonucleoprotein (RNP) complexes. Additionally, DDX3 is also involved in pathogen sensing and interferon (IFN) activation [39,40]. UBR5 is a member of the HECT E3 ubiquitin ligase family, whose functions remain incompletely understood. Only a subset of its known interaction partners are targeted for ubiquitination [38]. Besides its role in transcription and translation, UBR5 also regulates the DNA damage response and promotes cell cycle progression [38].

Considering that m139 can form a complex with m140 and m141 in MCMV infected cells [24], we tested whether m139 can interact with DDX3 or UBR5 in the absence of m140 and m141. For this, DDX3 or UBR5 were immunoprecipitated from lysates of HEK-239A cells transfected with HA-tagged versions of m139, m140, or m141. Under these conditions, only m139 co-precipitated with DDX3 and UBR5 (Figure 3B and C). This suggests that m139 itself specifically interacts with DDX3 and UBR5 and that m140 and m141 are not required for these interactions.

Next, we tested whether m139 affects the subcellular distribution of DDX3 and UBR5 in infected cells. To do this, we infected SVEC4-10 endothelial cells with MCMV m139-HA or MCMV m139*stop*. DDX3 showed a predominantly cytoplasmic distribution in uninfected cells. In MCMV-infected cells it also accumulated in nuclear replication compartments, and this re-distribution was dependent on m139 expression (Figure 4A). UBR5 was detected only in the nucleus. It also accumulated in nuclear replication compartments of infected cells, but this re-distribution was not dependent on m139 expression (Figure 4B), suggesting that the accumulation of UBR5 in viral replication compartments occurs by other means.

**Figure 4.**
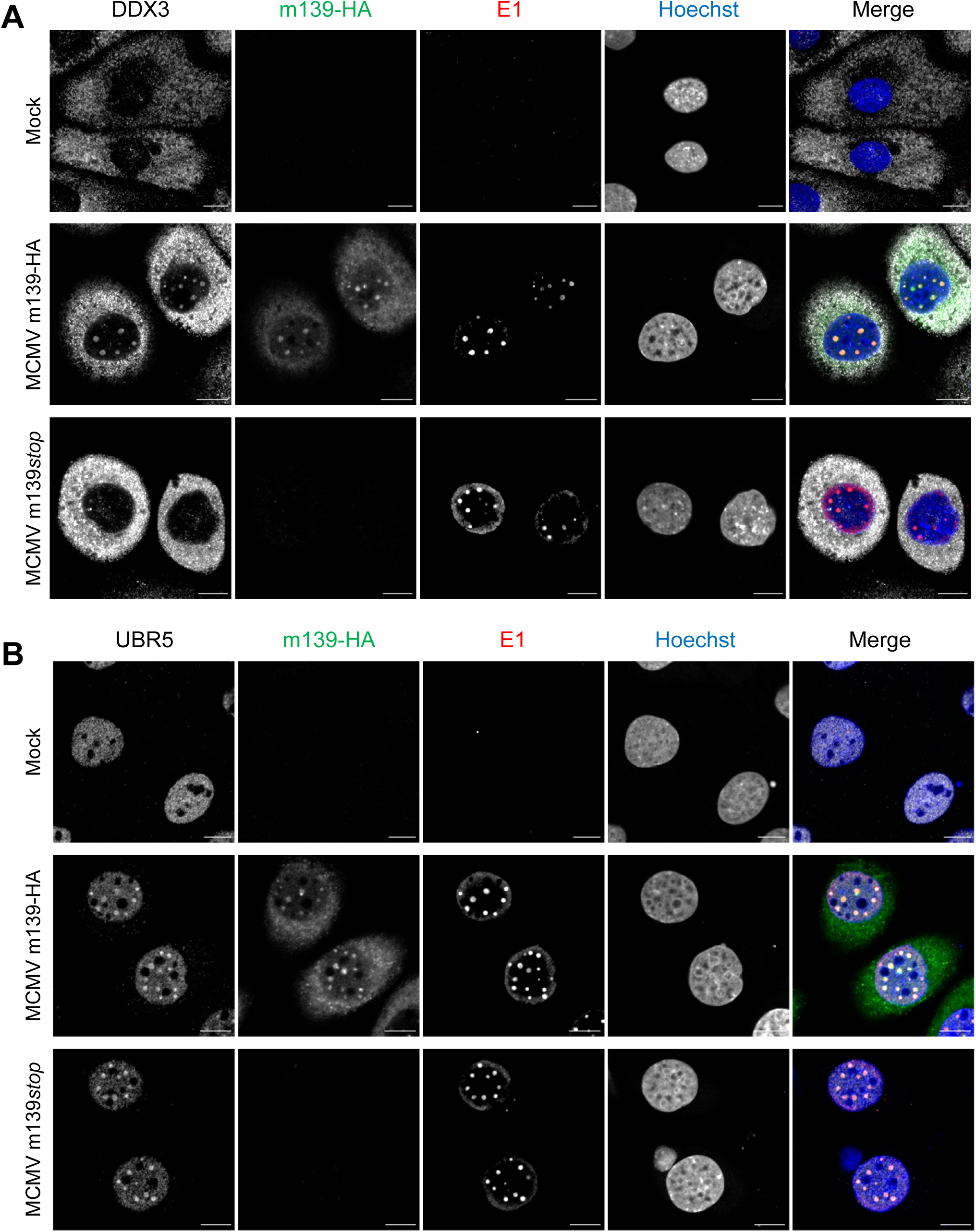
DDX3 and UBR5 are recruited to the viral replication compartments upon MCMV infection. SVEC4-10 endothelial cells were infected with MCMV m139-HA or MCMV m139*stop* (MOI=1) and fixed at 8 hpi. The subcellular localization of DDX3 (A), UBR5 (B), and the viral m139 and E1 (M112-113) proteins was analyzed by immunofluorescence using protein-specific and anti-HA antibodies. Nuclei were counterstained with Hoechst 33342. Representative images acquired by confocal microscopy are shown. Scale bar, 10 µm.

### DDX3 and UBR5 restrict MCMV replication in SVEC4-10 endothelial cells

The DEAD box RNA helicase DDX3 acts as a proviral factor for several RNA viruses, but also participates in the innate immune response, thereby functioning as an antiviral factor [39,40]. As MCMV replication in SVEC4-10 endothelial cells and iBMDM was impaired in the absence of m139 (Figure 2C and D), we wanted to determine whether DDX3 or UBR5 are responsible for the impaired replication of the m139*stop* mutant. Therefore, we generated DDX3 and UBR5-deficient SVEC4-10 cells by CRISPR/Cas9 gene editing. Despite great efforts we were unable to obtain completely DDX3-deficient cells. However, we were able to isolate a SVEC4-10 cell clone that expresses DDX3 in very small amounts (Figure 5A). In contrast, it was not difficult to isolate UBR5-deficient SVEC4-10 cells (Figure 5B). Remarkably, replication of MCMV m139*stop* could be rescued to WT levels in DDX3-deficient (Figure 5C) as well as in UBR5-deficient cells (Figure 5D). These findings suggested that DDX3 and UBR5 act as restriction factors of MCMV replication and that m139 counteracts this restriction.

**Figure 5.**
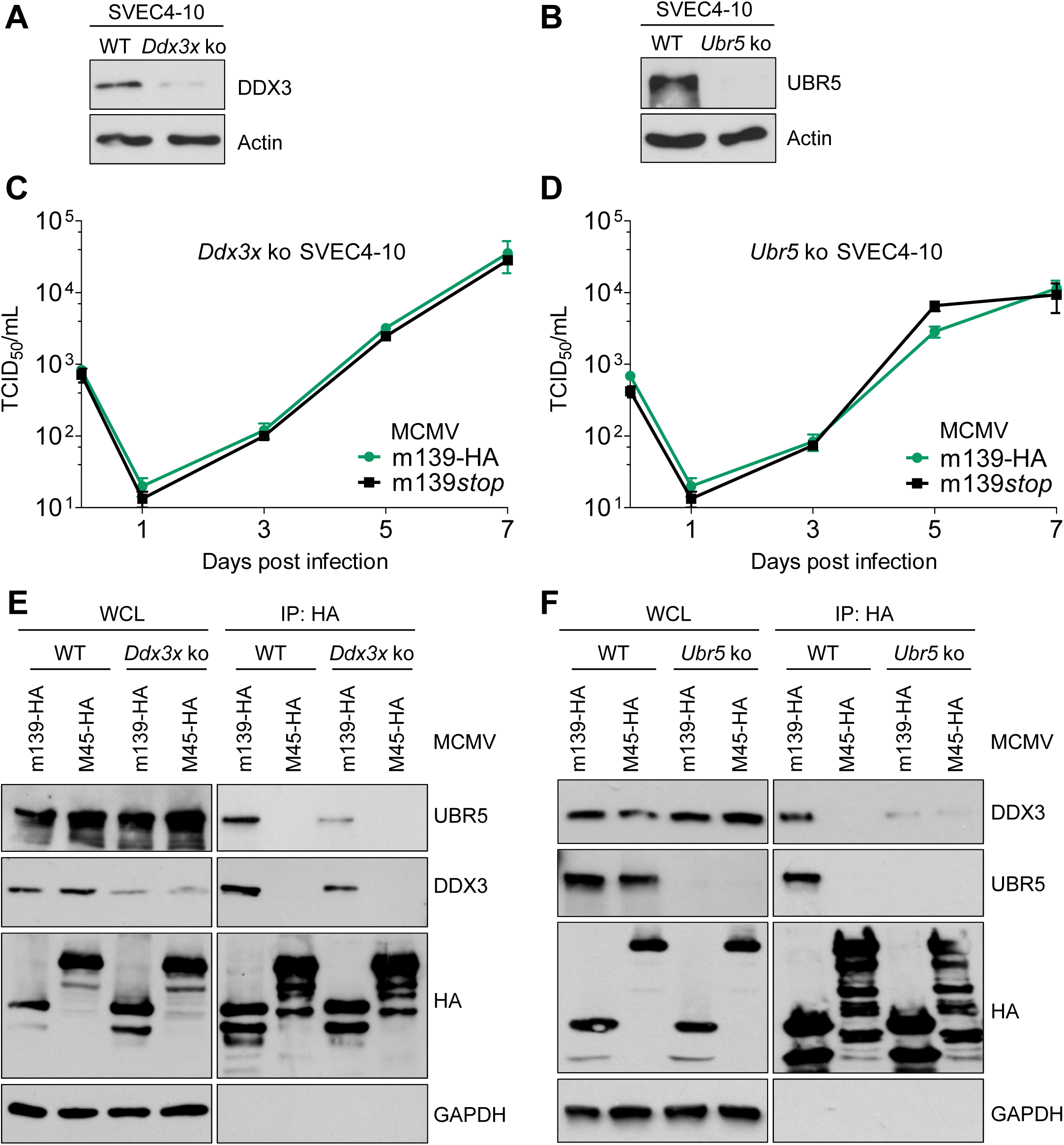
Replication defect of MCMV m139*stop* is rescued in DDX3- and in UBR5-deficient SVEC4-10 cells. (A, B) *Ddx3x* and *Ubr5* ko SVEC4-10 cells were generated by CRISPR/Cas9 gene editing. DDX3 (A) and UBR5 (B) expression was verified by immunoblot analysis. Note that the *Ddx3x* ko was incomplete. (C, D) For multistep replication kinetics, *Ddx3x* (C) and *Ubr5* (D) ko SVEC4-10 cells were infected with MCMV m139-HA and MCMV m139*stop* at an MOI of 0.01. At different times points post infection, supernatants were collected for titration. Viral titers are shown as means ±SEM. (E, F). WT and *Ddx3x* ko (E) or Ubr5 ko (F) SVEC4-10 cells were infected with MCMV m139-HA or MCMV M45-HA (control) at an MOI of 5 and harvested 24 hpi. The m139 protein was immunoprecipitated using an anti-HA affinity matrix. Co-precipitating proteins were detected by immunoblotting.

As both, DDX3 and UBR5, were involved in restraining the replication of MCMV m139*stop*, we interrogated whether the interaction of m139 with these two proteins is interdependent. We tested this in immunoprecipitation experiments with DDX3 and UBR5-deficient cells. Indeed, reduced amounts of UBR5 co-precipitated with m139 in DDX3-deficient cells (Figure 5E), and reduced amounts of DDX3 co-precipitated with m139 in UBR5-deficient cells (Figure 5F). These results indicate that the interaction of m139 with UBR5 is dependent on DDX3 and vice versa. It is conceivable that m139 forms a heterotrimeric complex with DDX3 and UBR5 within viral replication compartments.

Considering that UBR5 is an E3 ubiquitin ligase that can mark interacting proteins for proteasomal degradation, we tested whether m139 induces DDX3 degradation by recruiting UBR5. We infected SVEC40-10 and iBMDM with MCMV m139-HA and m139*stop* and analyzed DDX3 levels over time. As shown in Figure S2, m139 expression by MCMV did not induce decreased DDX3 levels. Thus, the recruitment of UBR5 by m139 does not seem to induce DDX3 degradation.

### m139 inhibits DDX3-mediated IFN-β induction

DDX3 is known to function as a mediator of the IFN-β antiviral response [39,40]. Moreover, a recent transfection-based screen for MCMV-encoded inhibitors of IFN-β induction identified m139 as one of several candidates [27]. To examine the effect of m139 on IFN-β induction we used immortalized bone marrow-derived macrophages expressing firefly luciferase under control of the endogenous IFN-β promoter (iBMDM β-luc). Infection of these cells with MCMV m139-HA resulted in an increase in luciferase expression, confirming that macrophages activate the IFN-β promoter upon MCMV infection [27,41]. Infection with MCMV m139*stop* resulted in significantly increased IFN-β promoter activation, suggesting that m139 dampens IFN-β induction upon MCMV infection (**Error! Reference source not found**.A).

Next, we analyzed IFN-β transcription by qRT-PCR upon MCMV infection of iBMDM macrophages and SVEC4-10 endothelial cells, the two cell types in which MCMV m139*stop* showed a replication defect (Figure 2). Infection with vesicular stomatitis virus (VSV), a very potent inducer of type I IFNs, was used as a positive control. As shown in Figure 6B, infection with MCMV m139*stop* resulted in significantly higher IFN-β transcription compared to cells infected with the m139-HA WT virus. By contrast, MCMV m139*stop* did not induce increased IFN-β transcription in SVEC4-10 endothelial cells (Figure 6C), suggesting that m139 curtails IFN-β induction in macrophages, but not in endothelial cells.

**Figure 6.**
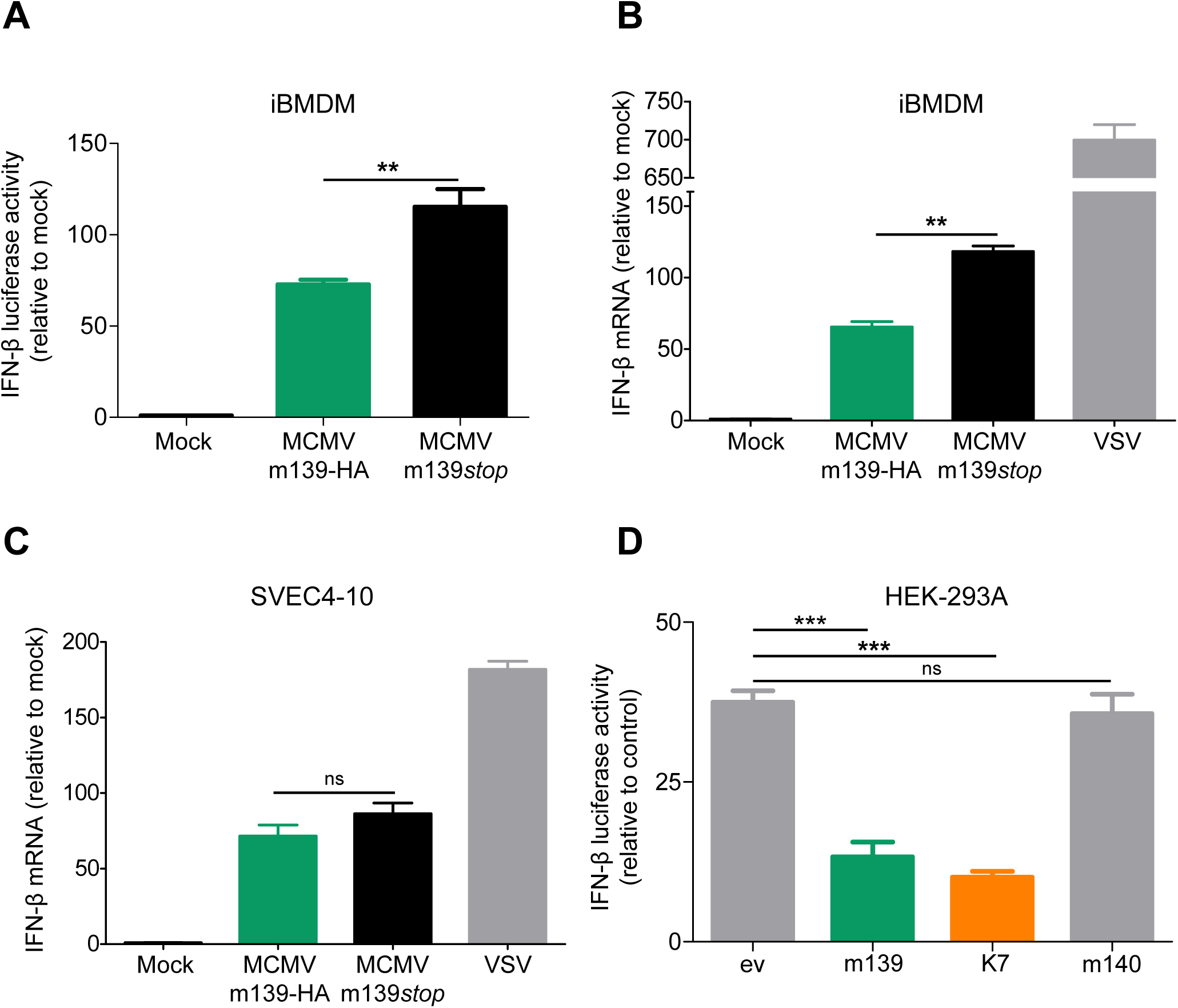
m139 curtails IFN-β transcription in macrophages. (A) iBMDM macrophages expressing firefly luciferase under the control of the endogenous IFN-β promoter were mock-infected or infected with MCMV m139-HA or m139*stop*. At 8 hpi, cells were lysed and luciferase activity was measured. Means ±SEM of 3 biological replicates are shown. (B, C) iBMDM (B) or SVEC4-10 (C) cells were infected with MCMV m139-HA, MCMV m139*stop* or vesicular stomatitis virus (VSV), a very potent IFN-β inducer. Cells were harvested 6 hpi for RNA isolation and qRT-PCR was performed for IFN-β. Data are shown as means ± SD of three biological replicates. (D) HEK-293A cells were co-transfected with expression plasmids for DDX3 and IKKε, a reporter plasmid containing firefly luciferase under the control of the murine IFN-β promoter (IFNβ-luc), a renilla luciferase normalization control, and an empty vector (pcDNA) or plasmids expressing MCMV m139, MCMV m140 as a negative control or VACV K7 as a positive control. Firefly luciferase activities were normalized to renilla luciferase activities from the same samples. Values were normalized to those of cells transfected with the luciferase vectors and empty vector, but without DDX3 and IKKε plasmids. Means ±SEM of 3 biological replicates are shown. Significance was determined by ANOVA. ns, not significant; **, p<0.01; ***, p<0.001.

The vaccinia virus (VACV) K7 protein has been shown to interact with DDX3 and block DDX3-mediated IFN-β induction by inhibiting IRF3 activation by the TBK1-IKKε complex [42]. To determine whether m139 inhibits DDX3-mediated IFN-β induction in a similar way as K7, we used a previously described luciferase reporter assay [42]. HEK-293A cells were transfected with DDX3 and IKKε expression plasmids to activate the IFN-β promoter by overexpression. Plasmids expressing m139, m140, or K7 were co-transfected, and IFN-β promoter activation was measured using a reporter plasmid expressing firefly luciferase under the control of the murine IFN-β promoter (IFNβ-Luc). In this assay, m139 and K7 strongly inhibited IFN-β promoter activation, whereas m140 did not (Figure 6D). These results confirmed that m139 inhibits DDX3-dependent IFN-β induction.

To determine whether the replication defect of MCMV m139*stop* in macrophages is dependent on DDX3-mediated IFN-β induction, we constructed a recombinant MCMV expressing K7 in place of m139 (Figure 7A). K7 expression by the K7[Δm139] substitution mutant was confirmed by immunoblotting (Figure 7B). Next, we assessed the replication properties of the substitution mutant in iBMDM and SVEC4-10 cells. In iBMDM, MCMV K7[Δm139] replicated to similar titers as the control virus (Figure 7C), indicating that the VACV K7 protein can rescue the replication defect of an m139-deficient virus in macrophages. By contrast, the K7[Δm139] substitution did not rescue the replication defect in SVEC4-10 endothelial cells (Figure 7D). This finding was not surprising as IFN-β transcription was not increased in the absence of m139 in endothelial cells (Figure 6C). Moreover, the replication defect of MCMV m139*stop* in SVEC4-10 cells depended on both DDX3 and UBR5 (Figure 5), which suggests that m139 antagonizes an IFN-independent function of DDX3 in endothelial cells.

**Figure 7.**
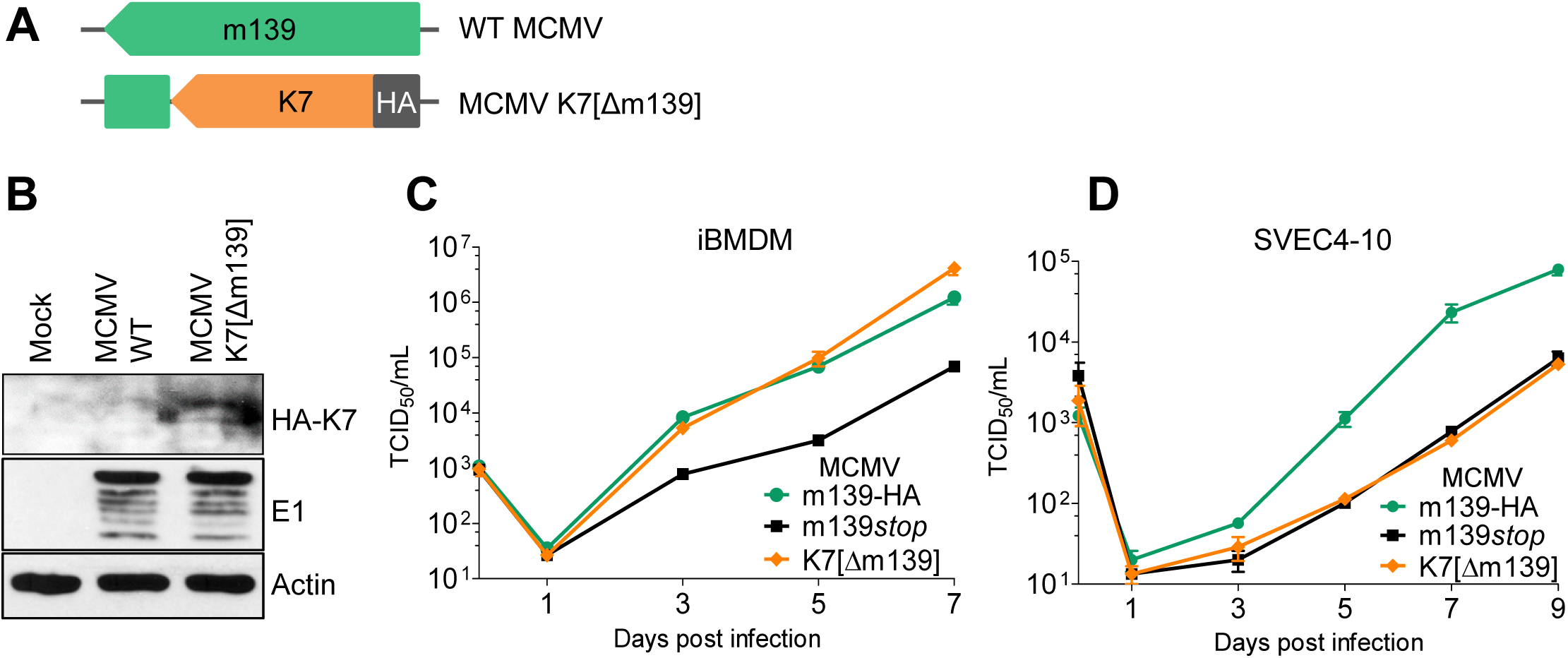
VACV protein K7 rescues the replication defect of an m139-deficient MCMV in macrophages. (A) Schematic representation of WT MCMV and MCMV K7[Δm139], a substitution mutant expressing HA-tagged VACV K7 instead of m139. (B) 10.1 cells were infected at an MOI of 5 with WT MCMV or K7[Δm139]. Cell lysates were collected 24 hpi and subjected to immunoblot analysis. (C) iBMDM were infected with MCMV m139-HA, m139*stop*, or K7[Δm139] at an MOI of 0.025. Virus release into the supernatant was quantified by titration. Viral titers are shown as means ±SEM of 3 biological replicates. (D) SVEC4-10 endothelial cells were infected at an MOI of 0.01 with the same viruses as in C.

### m139 is crucial for viral dissemination in vivo

To analyze the importance of m139 for MCMV dissemination in its host, we chose a route of infection that strongly depends on macrophages. MCMV administered by footpad-inoculation rapidly reaches the popliteal lymph nodes (PLN), where it infects subcapsular sinus macrophages. From there it spreads slowly to distant sites, suggesting that lymph node macrophages can be a bottleneck to viral dissemination [43]. We infected BALB/c mice with MCMV m139-HA, m139*stop*, or K7[Δm139] and measured viral titers in PLN on day 3 post infection. Compared to the HA-tagged WT virus (MCMV m139-HA), infection with MCMV m139*stop* resulted in strongly reduced viral titers in PLN (Figure 8A). Expression of K7 by the K7[Δm139] substitution mutant did not restore viral titers in PLN to WT levels, but moderately improved titers compared to the m139*stop* mutant. Hence, m139 is required for efficient spread to or replication in PLN.

**Figure 8.**
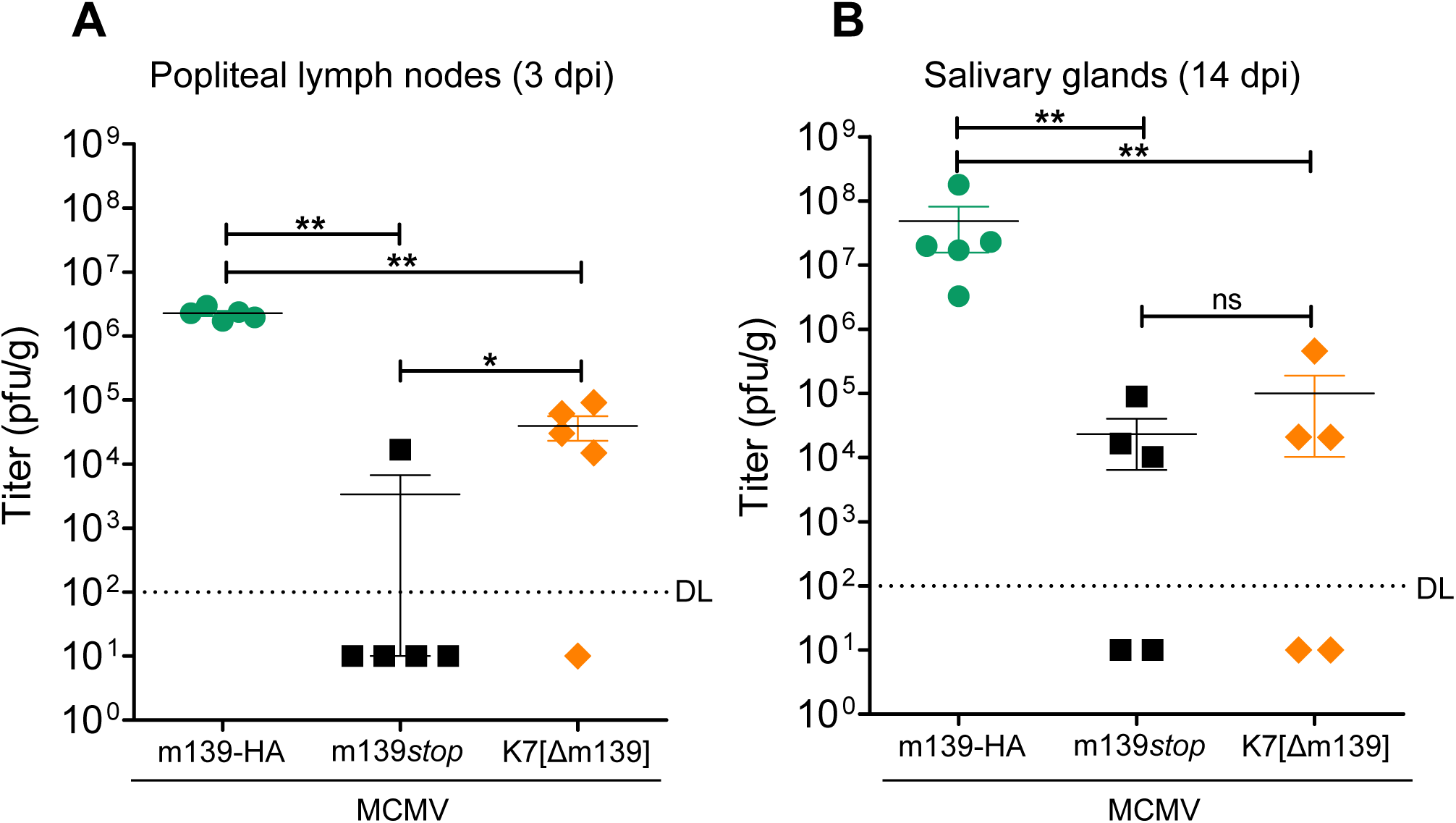
m139 is crucial for MCMV dissemination to popliteal lymph nodes and salivary glands. BALB/c mice were infected by injection of 10^5^ pfu MCMV m139-HA, m139*stop*, or K7[Δm139] into the footpad. (A) Popliteal lymph nodes (PLN) were harvested 3 days post infection. (B) Salivary glands were harvested 14 days post infection. Viral titers were determined by plaque assay. Mean titers ±SEM are shown. DL, detection limit; ns, not significant; *, p<0.05; **, p<0.01.

Next, we analyzed MCMV dissemination to the salivary glands (SG), which are important organs for viral persistence and shedding [44]. Viral titers in SG homogenates were determined on day 14 post infection. Titers in salivary glands of mice infected with MCMV m139*stop* and K7[Δm139] MCMV were strongly reduced (Figure 8B), which confirms the importance of m139 for viral dissemination *in vivo*. The substitution of m139 by K7 in the K7[Δm139] recombinant virus did not rescue the viral dissemination defect of the m139-deficient virus. Similar to MCMV m139*stop*, K7[Δm139] has a replication defect in endothelial cells (Figure 8C) and possibly in further cell types, suggesting that the VACV DDX3 antagonist K7 does not fulfill all functions of m139, particularly those related to UBR5. In summary, the results show that m139 is required for efficient dissemination from a peripheral site of infection to local lymph nodes and to distant sites.

## Discussion

In this study, we show that MCMV m139 interacts with the host proteins DDX3 and UBR5 and inhibits DDX3-mediated induction of IFN-β in macrophages. DDX3 is known to play an important role in antiviral innate immunity [39,40]. It participates in RNA sensing by RIG-I and related RNA helicases, interacts with the kinases TBK1 and IKKε to activate IRF3, and can bind directly to the IFN promoter [39,40]. More recently, DDX3 was also shown to interact with IKKα and facilitate IRF7 activation by NIK and IKKα [45]. Two known sensors of MCMV infection, ZBP1/DAI and TLR9 [46-49], can induce type I IFNs through these kinase complexes [32,45]. Interestingly, IRF7 is expressed only in lymphocytes and myeloid cells, which comprise monocytes, macrophages, and dendritic cells [50]. This could be an explanation for the increased IFN-β transcription upon infection with MCMV m139*stop* in macrophages but not in endothelial cells (Figure 6B and C). Another possible reason could be the expression of the relevant pattern recognition receptors. TLR9, for instance, is expressed predominantly by antigen presenting cells such as macrophages and dendritic cells [51]. Only a few viral antagonists of DDX3-mediated IFN signaling have been described: The influenza virus protein PB1-F2 inhibits DDX3 by inducing its proteasomal degradation, while VACV protein K7 and hepatitis B virus polymerase block the interaction of DDX3 with IKKε [42,52,53]. This study describes the first DDX3 antagonist encoded by a herpesvirus, which inhibits IFN production in a similar fashion as VACV K7 (Figure 9A).

**Figure 9.**
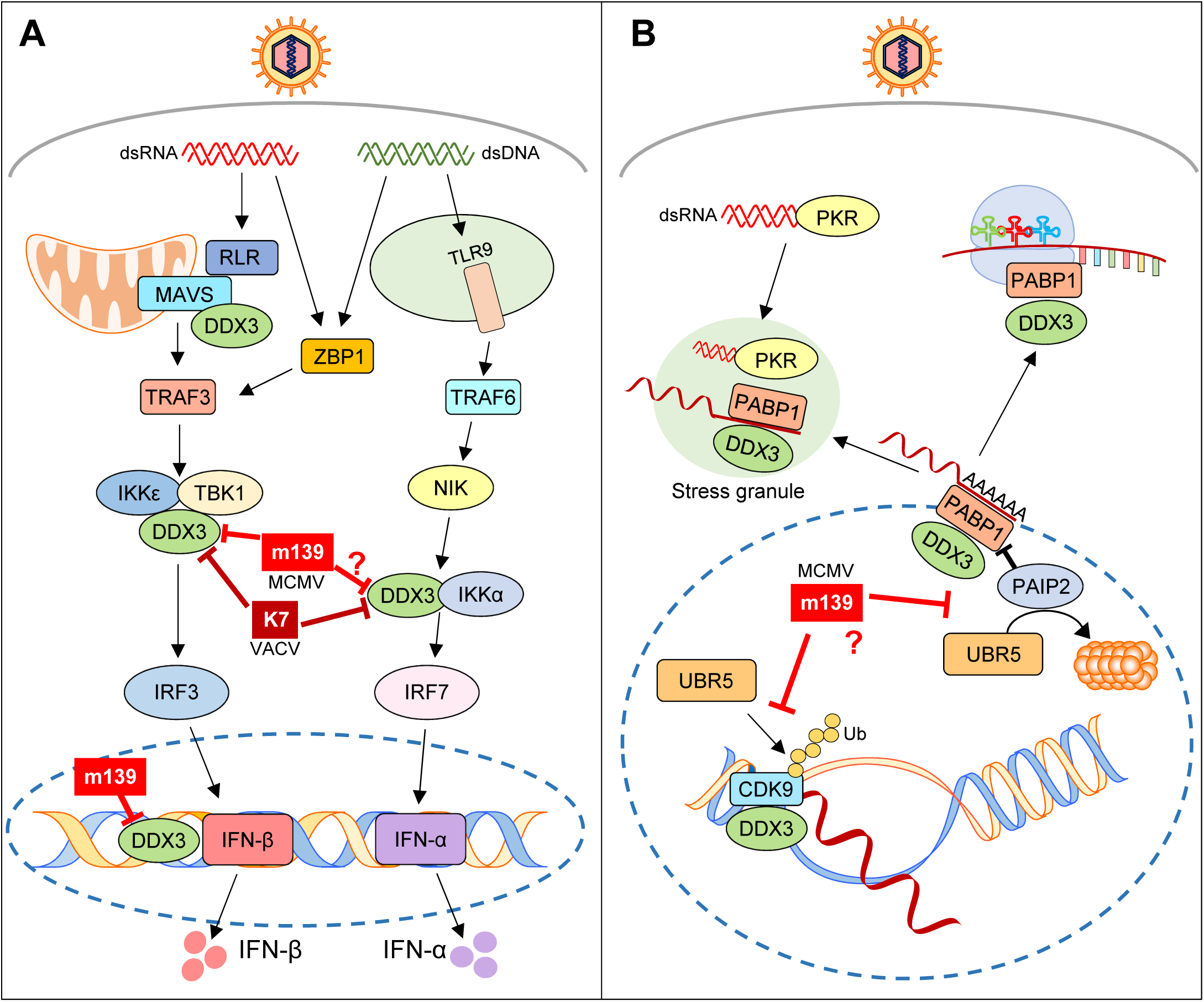
Known functions of DDX3 and UBR5 and their presumed targeting by m139. (A) Functions of DDX3 in IFN activating pathways. MCMV m139 interacts with DDX3 and inhibits IFN-β transcription either by targeting the interaction of DDX3 with IKKε (similar to VACV K7) or by blocking DDX3 binding to the IFN-β promoter; m139 might also inhibit IFN-α induction in same way as K7. (B) Functions of DDX3 and UBR5 in RNA transcription, processing, and translation. MCMV m139 co-localizes with DDX3 and UBR5 in the nucleus, suggesting that m139 antagonizes DDX3 and UBR5-dependent functions in the nucleus.

Besides its role in innate immunity, DDX3 also participates in several aspects of RNA processing, including mRNA transcription, splicing, export, and translation initiation [54]. These functions affect a subset of cellular and viral RNAs containing specific structural features. For instance, DDX3 affects cell cycle regulation by facilitating nuclear export and translation of several cyclin mRNAs [54]. DDX3 also facilitates Rev-dependent nuclear export and translation of HIV mRNAs, demonstrating that viruses can exploit DDX3 for their own ends [39,54]. Although much less is known about the specific functions of UBR5, it is interesting to note that UBR5 is also involved in the regulation of transcription and translation [38]. While DDX3 promotes translation of specific mRNAs by interacting with the translation initiation factors eIF4A, eIF4G, and the polyadenylate-binding protein PABP1 [54], UBR5 supports this process by inducing the degradation of the PABP-interacting protein 2 (PAIP2), which functions as an inhibitor of PABP1 [55]. The fact that MCMV m139 interacts with both, DDX3 and UBR5 (Figure 3A-C), and co-localizes with both proteins in viral replication compartments (Figure 4) suggests that m139 might regulate the transcription, processing, or translation of specific viral or cellular mRNAs (Figure 9B) that affect the efficiency of viral replication in endothelial cells.

The m139 gene is a member of the US22 gene family, which has 12 members in MCMV and 12 members in HCMV. Therefore, it would be interesting to know if m139 has a functional homolog in HCMV. Based on phylogenetic analysis, the closest sequence homologs are the HCMV genes US22, US23, US26. Little is known about the function of the corresponding proteins, but a proteomic study has identified the US22 protein as an RNA-binding protein [35]. The same study also found increased amounts of DDX3 bound to mRNAs of HCMV-infected cells, but it remains unknown whether or not these two observations are functionally connected. Another study has shown by siRNA-mediated knockdown that DDX3 acts as a proviral host factor during HCMV infection [34]. A recent mass spectrometry-based interactome analysis of all HCMV proteins did not identify DDX3 as a high-confidence interactor, but UBR5 was found to interact with six viral proteins, two of which are members of the US22 family: US26 and UL36 [56]. Moreover, another study demonstrated a role of PAIP2, whose stability is regulated by UBR5/EDD1, as a restriction factor limiting productive HCMV infection [33].

Small animal models, such as the mouse model of CMV infection, help to verify the biological significance of identified virus-host interactions in an organism. Under ideal circumstances, the loss of a viral protein can be compensated by using a mouse strain in which the target protein has been removed by genetic knockout. This is promising if the viral protein only targets one host protein (e.g. [16,57]). In the present case, such a genetic rescue is unlikely to work as m139 interacts with at least two host proteins, DDX3 and UBR5. Nevertheless, a knockout of DDX3 or UBR5 in cultured SVEC4-10 endothelial cells was sufficient to restore the replication of the MCMV m139*stop* mutant to WT levels (Figure 5C and D), suggesting that DDX3 and UBR5 impinge on the same pathway. Unfortunately, the same experiment cannot be done in mice as *Ddx3x* and *Ubr5* knockout mice are not viable [58,59]. One way to overcome this obstacle is the use of conditional knockouts such as the recently published conditional *Ddx3x* knockout mice [60]. Another approach consists of replacing the viral gene by another gene of known function, as we have done it by replacing MCMV m139 with a known DDX3 antagonist, VACV K7. Indeed, the substitution mutant rescued MCMV replication in BMDM in cell culture (Figure 7C). However, the substitution provided only a partial rescue in vivo (Figure 8A), probably because K7 cannot compensate for all functions of m139, particularly those related to UBR5.

Obviously, it would be highly interesting to know which DDX3 and UBR5-dependent functions m139 antagonizes in the nucleus of SVEC4-10 endothelial cells. Both, DDX3 and UBR5 affect the transcription and translation initiation of a subset of mRNAs (Figure 9B). Only a few mRNAs transcripts are known targets. However, the exact structural requirements for DDX3-mediated regulation and the complete set of transcripts affected are currently unknown. Additional research, which is beyond the scope of the present study, will be necessary to understand the details of transcriptional and translational regulation by DDX3 and UBR5 and to clarify how viral interference promotes viral replication.

## Materials & Methods

### Ethics statement

Animal experiments were performed according to the recommendations and guidelines of the FELASA (Federation for Laboratory Animal Science Associations) and Society of Laboratory Animals (GV-SOLAS) and approved by the institutional review board and local authorities (Behörde für Gesundheit und Verbraucherschutz, Amt für Verbraucherschutz, Freie und Hansestadt Hamburg, reference number 017/2019).

### In vivo experiments

Six to eight-week-old BALB/c female mice (Janvier Laboratories) were infected by injecting 10^5^ PFU of MCMV into the left footpad. PLN were harvested on day 3 and SG on day 14 post infection. Organ homogenates were titrated by plaque assay on M2-10B4 cells essentially as described [44].

### Cell lines

SVEC4-10 (CRL-1658), NIH-3T3 (CRL-1658), TCMK-1 (CCL-139), M2-10B4 (CRL-1972), and HEK 293T (CL-11268) were obtained from the ATCC. HEK-293A cells (R705-07) were purchased from Invitrogen. Murine 10.1 fibroblasts have been described [61]. Immortalized murine bone marrow-derived macrophages (iBMDM) were obtained from BEI Resources, NIAID NIH (NR-9456). BMDM expressing firefly luciferase under control of the endogenous IFN-β promoter were isolated from transgenic mice [62] and immortalized by transduction with retrovirus J2 as described [63]. All cells were cultured at 37°C and 5% CO_2_ in complete Dulbecco’s modified Eagle medium (DMEM) supplemented with 10% fetal calf serum and 100 IU penicillin/100 μg streptomycin.

### Viruses

The repaired WT MCMV Smith bacterial artificial chromosome (BAC), pSM3fr-MCK-2fl [64], was used to construct mutant MCMVs by *en passant* mutagenesis [65]. The m139-HA mutant was generated by inserting the hemagglutinin (HA) epitope sequence at the 3’ end of the m139 ORF. m139*stop* was generated based on an HA-tagged mutant by introducing a point mutation at the position 195923 (T→A), which resulted in a stop codon at position 32 of m139. To generate MCMV K7[Δm139], the HA-K7 sequence was PCR-amplified from pCMV-HA-K7R and used to replace m139. To exclude unintended mutations, all recombinant BACs were sequenced at the NGS facility of the Heinrich Pette Institute. Infectious MCMV was reconstituted by transfection of 10.1 cells with BAC DNA. Virus stocks were titrated using the median tissue culture infective dose (TCID_50_) method. Centrifugal enhancement (1000 × *g*, 30 min) was applied for high-MOI infections. VSV-GFP [66] was kindly provided by César Muñoz-Fontela (Bernhard Nocht Institute for Tropical Medicine, Hamburg, Germany).

### Growth curves

Multistep replication kinetics were done as described [61]. Cells were seeded and infected in 6-well dishes. Three hours post infection cells were washed twice with phosphate-buffered saline (PBS), and fresh medium was added. Supernatants harvested from infected cells were titrated on 10.1 cells.

### Plasmids

pcDNA3-HA-m139, pcDNA3-HA-m140, and pcDNA-m141-HA were generated by PCR-amplification of the respective ORF from the MCMV BAC with primers containing restriction sites and an HA tag sequence. The PCR products were cloned in pcDNA3 (Invitrogen) using EcoRI and EcoRV restriction sites HA-m139, HindIII and XbaI for HA-m140, and HindIII and XbaI for m141-HA. pCMV-HA-K7R [42] was provided by Martina Schröder (Maynooth University, Maynooth, Ireland). pcDNA3-IKKε, encoding human I-kappa-B kinase epsilon [67], was provided by Rongtuan Lin (McGill University, Montreal, Canada). pcDNA-HA-DDX3 was obtained from Addgene (plasmid 44975). pGL3basic-IFNβ-Luc (IFNβ-Luc) has been described [41]. pRL-TK was purchased from Promega. Transfection of NIH-3T3 fibroblasts was done using lipofectamine (Thermo Fisher Scientific) according to the manufacturer’s instructions.

### Generation of knockout cells

The lentiviral CRISPR/Cas9 vector pSicoR-CRISPR-PuroR was used was used for generation of *Ddx3x* and *Ubr5* knockout SVEC4-10 cells essentially as previously described [68]. Guide RNAs targeting *Ubr5* (GGCTACTATTAAACAGTGTG) and *Ddx3x* (CTATCTTTACTAGAACTCCA) were designed using E-CRISP (http://www.e-crisp.org/E-CRISP). Lentivirus production, transduction of SVEC4-10 cells, and selection of clones was done as previously described [68]. Cell clones were screened by immunoblot analysis.

### Antibodies and reagents

Monoclonal antibodies against HA (3F10; Roche), β-actin (AC-74; Sigma), glyceraldehyde-3-phosphate dehydrogenase (GAPDH) (14C10; Cell Signaling), DDX3 (C-4; Santa Cruz Biotechnology), UBR5 (B-11; Santa Cruz Biotechnology) and polyclonal antibodies against LSD1 (Cell Signalling) were obtained from commercial sources. Antibodies against MCMV IE1 (CROMA101) and M112-113 (CROMA103) were provided by Stipan Jonjic (University of Rijeka, Rijeka, Croatia). Secondary antibodies coupled to horseradish peroxidase (HRP) were purchased from DakoCytomation or Jackson ImmunoResearch. Secondary antibodies coupled to Alexa-488, Alexa-555 or Alexa-633 were purchased from Thermo Fisher Scientific.

### Immunofluorescence

Cells were seeded on 8-well µ-slides (Ibidi) one day before transfection or infection. On the following day, cells were transfected with expression plasmids using lipofectamine transfection reagent (Thermo Fisher Scientific), or infected at an MOI of 1 TCID_50_/cell. At the defined time points, cells were fixed using 4% Paraformaldehyde for 15 minutes at RT. The remaining aldehyde groups were blocked using 50 mM ammonium chloride. Cells were permeabilized with 0.3% Triton X-100 for 15 min and blocked with 0.2% gelatin (Sigma). Antibodies were diluted in 0.2% gelatin, applied to the cells, and incubated for 1 hour. Hoechst 33342 (Thermo Fisher Scientific) was used to stain nuclear DNA. Fluorescence images were acquired with a Nikon A1 confocal laser scanning microscope.

### Luciferase reporter assay

One day before infection, 3 × 10^5^ iBMDM IFN-β-luc reporter macrophages were seeded in 12-well plates. Cells were infected at an MOI of 3 TCID_50_/cell, washed 3 hpi with PBS, and incubated with fresh growth medium. 8 hpi cells were lysed in Cell Culture Lysis Reagent (Promega). Lysates were combined with Luciferase Substrate (Promega) and luminescence was measured with a FLUOstar Omega plate reader (BMG Labtech). For the Dual Luciferase Assay (Promega), 1.5 × 10^5^ HEK-293A cells were seeded in 12-wells plates and transfected using Lipofectamine 2000 (Thermo Fisher Scientific) with 180 ng pcDNA3-DDX3-HA, 180 ng pcDNA3-IKKε, 300 ng pGL3basic-IFNβ-Luc, 30 ng pRL-Renilla, and 500 ng of a viral protein expression plasmid or an empty pcDNA3 vector. 24 hours post transfection, cells were lysed and firefly and renilla luciferase activities were measured on a FLUOstar Omega reader (BMG Labtech) and evaluated according to the manufacturer’s protocol.

### Cell fractionation

Nuclear and cytoplasmic fractions of MCMV-infected 10.1 cells (2 × 10^6^ cells/well) were lysed stepwise using an NE-PER™ nuclear and cytoplasmic extraction kit (Thermo Fisher Scientific) according to the manufacturer’s instructions. Protein concentrations were determined using a BCA assay (Thermo Fisher Scientific). Equal quantities of nuclear and cytoplasmic proteins were separated by SDS-PAGE and analyzed by immunoblotting

### Immunoprecipitation and immunoblotting

For immunoprecipitation, cells were seeded in 6-well plates one day before infection. The following day the cells were infected at an MOI of 3 TCID_50_/cell. 24 hours post infection cells lysed using NP-40 buffer (50 mM Tris, 150 mM NaCl, 1% Nonidet P-40, and Complete Mini protease inhibitor cocktail [Roche]). Insoluble material were removed by centrifugation. The remaining supernatant was precleared with protein G Sepharose (PGS, GE Healthcare). m139-HA was immunoprecipitated with anti-HA Affinity Matrix (clone 3F10, Roche). DDX3 and UBR5 were precipitated with specific antibodies and PGS beads. Precipitates were washed 3 times with buffer 1 (1 mM Tris pH 7.6, 150 mM NaCl, 2 mM EDTA, 0.2% NP-40), twice with buffer 2 (1 mM Tris pH 7.6, 500 mM NaCl, 2mM EDTA, 0.2% NP-40) and once with buffer 3 (10mM Tris pH 7.6). They were then eluted by boiling in SDS-PAGE sample buffer (125 mM Tris pH 6.8, 4% SDS, 20% glycerol, 10% β-mercaptoethanol, 0.002% bromophenol blue) and subjected to SDS-PAGE and immunoblot analysis.

For immunoblot analysis, SDS-PAGE sample buffer or NP-40 buffer were used. Equal amounts of protein (NP-40) or lysate (SDS-PAGE sample buffer) were loaded to SDS-PAGE and subsequently transferred to a nitrocellulose (Amersham) or 0.2 µm polyvinylidene difluoride (Immobilon-PSQ, Sigma) by semi-dry or wet blotting. Proteins of interest were detected with protein-specific primary antibodies and HRP-coupled secondary antibodies by enhanced chemiluminescence (GE Healthcare) and imaged using X-ray films or with Fusion Capture Advance FX7 16.15 (Peqlab).

### SILAC, AP-MS, and LC-MS/MS

For SILAC, SVEC4-10 cells were grown for 8 passages in SILAC medium, supplemented with 10% dialyzed FCS, 4 mM glutamine, 100 IU penicillin/100 μg streptomycin in the presence of either ^13^C_6_,^15^N_2_-lysine / ^13^C_6_,^15^N_4_-arginine (SILAC heavy) or unlabeled lysine / arginine (SILAC light). Heavy and light labeled SVEC4-10 cells were infected with MCMV m139-HA and MCMV WT (MOI=5) and lysed 24 hours post infection in sterile-filtered NP-40 buffer. 1 mg of each whole cell lysate was used for immunoprecipitation with an anti-HA affinity matrix. Immunoprecipitates were washed six times with sterile-filtered minimal washing buffer (50 mM Tris, 150 mM NaCl, 10% (v/v) glycerol, pH 7.5) and eluted by boiling in elution buffer (1% (w/v) SDS in 50 mM Tris, 150 mM NaCl, pH 7.5). The eluted samples from heavy and light-labeled cells were mixed in a 1:1 ratio, separated by a short (1 cm) SDS-PAGE run to remove interfering substances, and stained with Coomassie R-250. Each lane was cut into 1 mm^3^ cubes followed by destaining. Disulfide bonds of proteins in the gel matrix were reduced in presence of 10 mM dithiotreitol (Fluka), alkylated in presence of 20 mM iodoacetamide (IAA, Sigma), and digested with trypsin (50:1 protein:enzyme ratio, Promega) overnight. Peptides were eluted from the gel pieces and dried in a vacuum concentrator. Samples were resuspended in 0.1% formic acid (FA) and transferred into a full recovery autosampler vial (Waters). Chromatographic separations were achieved on a Dionex Ultimate 3000 UPLC system (Thermo Fisher Scientific) with a two-buffer system (buffer A: 0.1% FA in water, buffer B: 0.1% FA in acetonitrile (ACN)). Attached to the UPLC was a C18 trapping column (Acclaim PepMap 100, 100 µm × 2 cm, 100 Å pore size, 5 µm particle size) for desalting and purification followed by a C18 analytical column (Acclaim PepMap 100, 75 µm × 50 cm, 100 Å pore size, 2 µm particle size). Peptides were separation using a 110 min gradient with increasing ACN concentration from 2 to 32% ACN. The eluting peptides were analyzed on a tribrid Quadrupole Iontrap Orbitrap mass spectrometer (Fusion, Thermo Fisher Scientific) in data-dependent acquisition (DDA) mode. For DDA, the mass spectrometer was operated in Orbitrap – Iontrap mode at top speed for precursor selection for fragmentation. Therefore, observed precursors with charge stages +2 to +5 in a range from 400 to 1300 m/z in a MS1 survey scan (2×10^5^ ions, 120,000 resolution, 120 ms fill time) were analyzed by MS/MS (HCD at 30 normalized collision energy, 1×10^4^ ions, 60 ms fill time) within 3 s. A dynamic precursor exclusion of 20 s was used.

### Data analysis and processing

Acquired DDA LC-MS/MS data were searched against the mouse SwissProt protein database downloaded from Uniprot (release August 2017, 16,909 protein entries) and an MCMV protein library using the Sequest algorithm integrated in the Proteome Discoverer software version 2.0. Mass tolerances for precursors was set to 10 ppm and 0.6 Da for fragments. Carbamidomethylation was set as a fixed modification for cysteine residues and ^13^C_6_,^15^N_2_ lysine, ^13^C_6_,^15^N_4_ arginine, the oxidation of methionine, pyro-glutamate formation at glutamine residues at the peptide N-terminus as well as acetylation of the protein N-terminus, methionine loss at the protein N-terminus and the acetylation after methionine loss at the protein N-terminus were allowed as variable modifications. Only peptide with a high confidence (false discovery rate < 1% using a decoy data base approach) were accepted as identified. Proteome Discoverer search results were imported into Skyline software version 4.2 allowing only high confidence peptides. Precursor traces (M, M+1, M+2) were extracted. Precursors with an idot product of >0.9 in at least one sample were kept. For each peptide, the ration of heavy to light was calculated and the median peptide ratio per protein was estimated, which were then used for relative comparison.

Amino acid sequence analysis of m139 and NES prediction was done using the computational tool Wregex [69].

### Quantitative RT-PCR

To quantify IFN-β transcripts, cells were infected at a MOI of 0.1 TCID_50_/cell. Three hpi cells were washed with PBS and fresh medium was added. Extraction of total RNA, cDNA synthesis and quantitative PCR (qPCR) was performed as described [68]. The following primers were used for amplification of transcripts: *Ifnb1* (CTGGCTTCCATCATGAACAA and AGAGGGCTGTGGTGGAGAA) and *Actb* (AGAGGGAAATCGTGCGTGAC and CAATAGTGATGACCTGGCCGT). *Ifnb1* transcripts were quantified using the ΔΔCt method and normalized to the housekeeping gene *Actb*.

### Statistical analysis

Statistical analyses were performed using GraphPad Prism 5.0 Software. One-way ANOVA with Bonferroni post test was used for the analysis of qRT-PCR and luciferase reporter assays. Statistical significance of in vivo experiments was assessed using the Mann-Whitney test.

## Acknowledgments

The authors would like to thank Martina Schröder, Rongtuan Lin, César Muñoz-Fontela, and Stipan Jonjic for providing materials and the NGS facility of the Heinrich Pette Institute (Daniela Indenbirken, Malik Alawi, and Sanamjeet Virdi) for sequencing.

## Data Availability

All relevant data are within the paper and its Supporting Information files.

---

**Figure S1.**
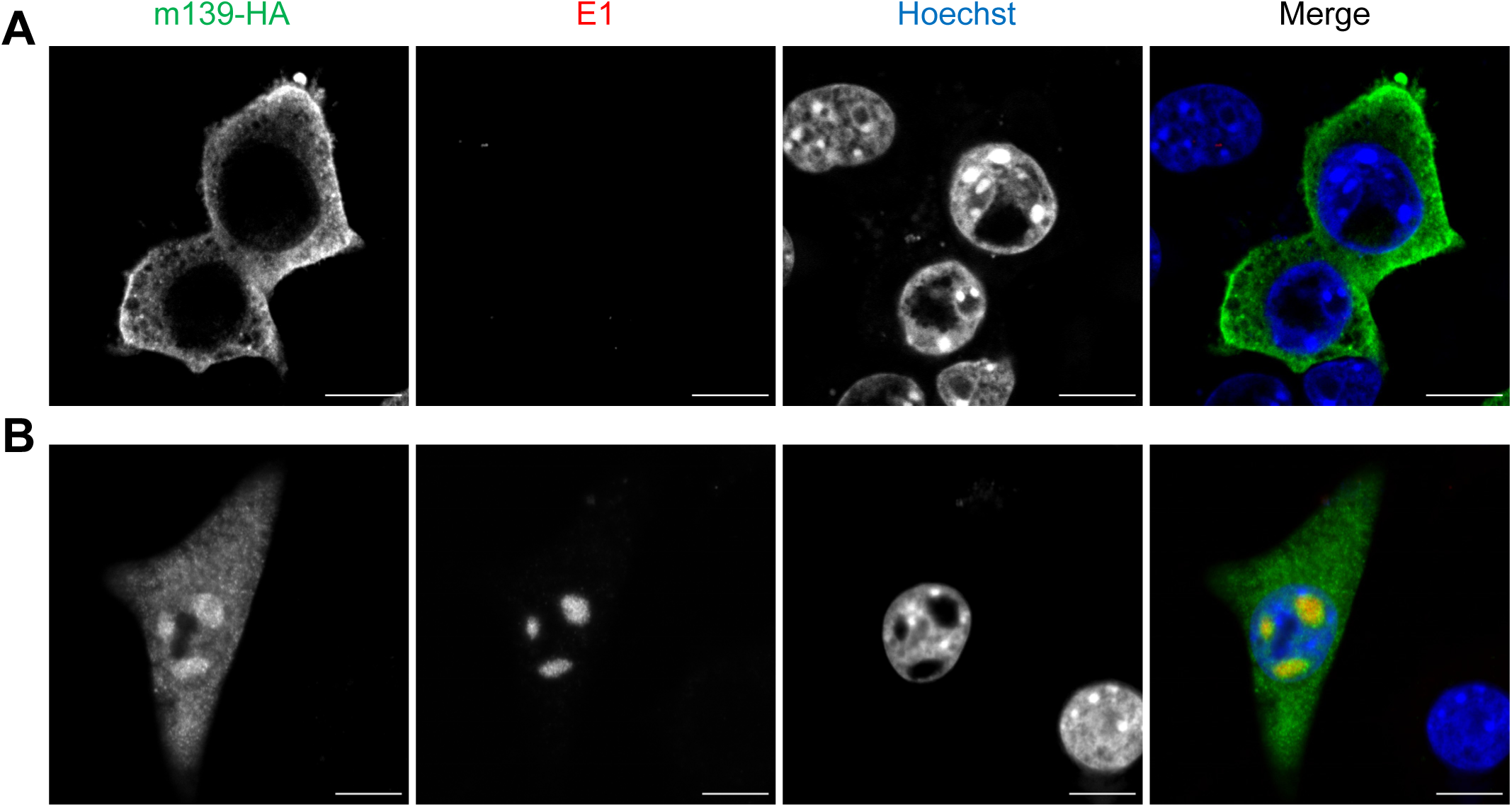
The m139 protein is recruited to the viral replication compartment by E1 proteins. NIH-3T3 fibroblasts were transfected with a plasmid encoding HA-tagged MCMV m139 (A) or plasmids encoding m139 and the MCMV E1 proteins (B). Cells were fixed 24 hpi and analyzed by immunofluorescence using antibodies specific for the HA epitope tag and E1. Nuclei were stained using Hoechst 33342. Representative images taken by confocal microscopy are shown. Scale bar, 10 µm.

**Figure S2.**
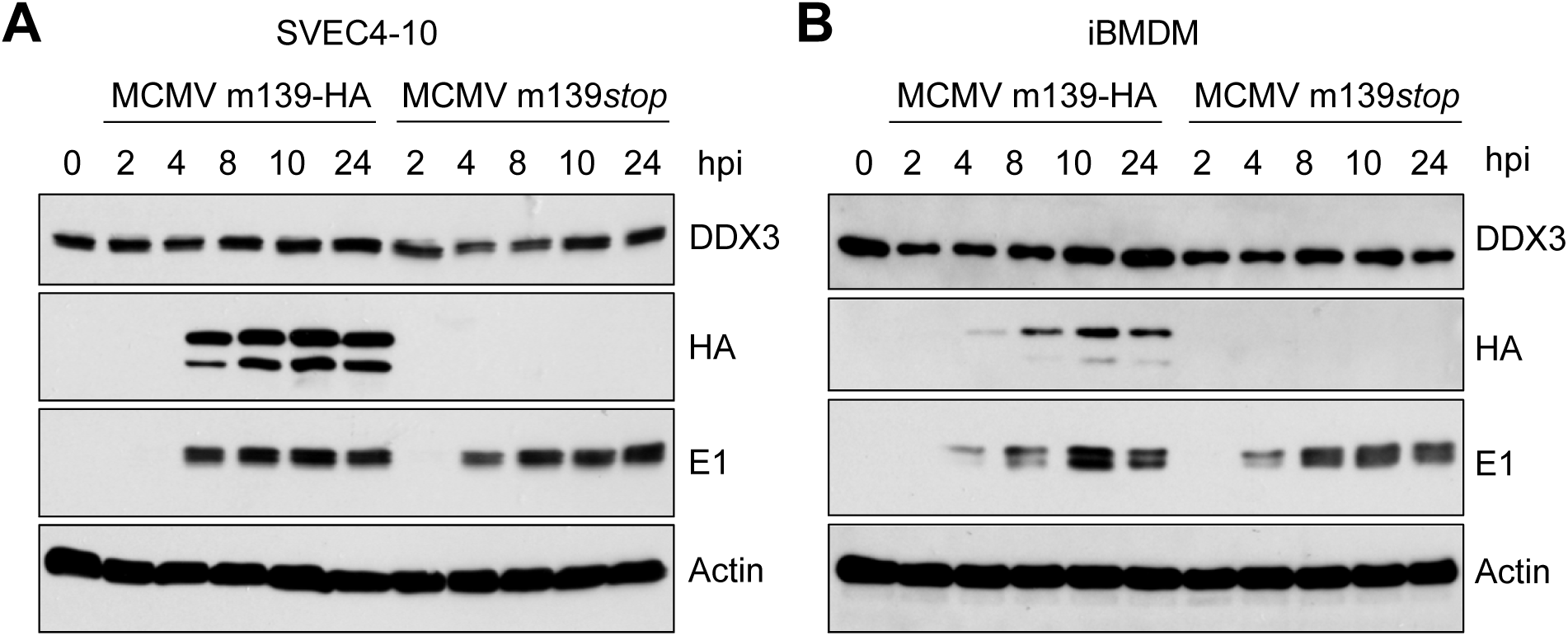
DDX3 levels are not decreased in MCMV-infected cells. (A) SVEC4-10 cells and (B) iBMDM were infected with MCMV m139-HA or MCMV m139*stop* at an MOI of 5. Whole cell lysates were prepared at the indicated times post infection and analyzed by immunoblot analysis.

